# Patterns of conservation and diversification in the fungal polarization network

**DOI:** 10.1101/154641

**Authors:** Eveline T. Diepeveen, Valérie Pourquié, Thies Gehrmann, Thomas Abeel, Liedewij Laan

## Abstract

The combined actions of proteins in networks underlie all fundamental cellular functions. Deeper insights into the dynamics of network composition across species and their functional consequences are crucial to fully understand protein network evolution. Large-scale comparative studies with high phylogenetic resolution are now feasible through the recent rise in available genomic datasets of both model *and* non-model species. Here we focus on the polarity network, which is universally essential for cell proliferation and studied in great detail in the model organism, *Saccharomyces cerevisiae*. We examine 34 proteins, directly related to cell polarization, across 200 fungal strains/species to determine evolution of the composition of the network and patterns of conservation and diversification.

We observe strong protein conservation for a group of 16 core proteins: 95% of all strains/species possess at least 75% of these proteins, albeit in varying compositions. We find high levels of variation in prevalence and sequence identity in the remaining 18 proteins, resulting in distinct lineage-specific compositions of the network in the majority of strains/species. We test if the observed diversification in network composition correlates with potential underlying factors and find that lineage, lifestyle, and genetic distance co-vary with network size. Yeast, filamentous and basal unicellular fungi form distinctive groups based on these analyses, with substantial differences to their polarization network. Our study shows that evolution of the fungal polarization network is highly dynamic, even between closely related species, and that functional conservation is achieved by varying structural conservation of the fungal polarization proteins.

**Data Deposition:** Sequence alignments used are available upon request.

## Introduction

All cells are maintained through fundamental cellular functions, such as respiration, biosynthesis, homeostasis and reproduction, that are crucial to the cell’s overall existence. These complex functions are carried out by the combined action of proteins in protein networks with distinct cellular tasks (Papin et al. 2005; Pawson & Nash 2003). Through evolution of protein networks, by means of e.g., amino acid mutations, network expansion/reduction, and interaction effects, diversity is generated that can ultimately lead to the evolution of new functions (see Gladieux et al. 2014 for a list of reviews). Protein networks can evolve by changes in network composition, changes in levels of protein conservation and divergence and changes in expression levels, each with potential functional consequences (Schüler & Bornberg-Bauer 2010; Voordeckers et al. 2015). Comparative genomics and/or interaction studies of protein networks, such as the citric-acid cycle (Huynen et al. 1999), mitotic checkpoint (Vleugel et al. 2012), and the mitogen-activated protein kinase pathway (Mody et al. 2009) illustrate such patterns. Most of these studies have examined overall protein network evolution by means of cross kingdom comparisons, covering ~20 divergent species. Although such approaches are insightful for testing if proteins are commonly found in distant clades of the tree of life, patterns such as parallel evolution are difficult to disentangle because of the lack of phylogenetic resolution. Examining (the factors promoting) protein network evolution in more species, especially at phylogenetic dense levels, is essential to gain deeper insights into the dynamics of protein network evolution.

What factors promote protein (network) evolution? Numerous factors have been presented that influence the evolution of individual proteins (see Pál et al. 2006; Zhang & Yang 2015). To simplify, these factors can be divided into two broad categories: sources of genetic variation, those relating to regional genomic properties, such as variation in mutation or recombination rate; and selection on genetic variation, factors dependent on specific protein properties, such as the proportion and distribution of sites that are involved in a specific functions, protein structure, expression level, and competition or adaptation (Pál et al. 2006). Although there are clear examples to illustrate these individual factors, the factors often do not act independently, making it hard to identify the relative importance of each factor. In yeast, for instance, the functional importance of a protein influences the rate of protein evolution (Wall et al. 2005; Drummond 2005; Hirsh & Fraser 2001), non-essential genes evolve on average faster than essential genes (Wall et al. 2005), and loci with more protein-protein interactions evolve on average slower (Jordan et al. 2003). Various studies have shown that expression rates have the most prominent effect on the rate of protein evolution (Wall et al. 2005; Drummond 2005).

Although selection acts on the outcome of fundamental cellular processes, specifically the phenotype, there are substantial differences in characteristics of proteins within a single network. As a consequence, selection can act differently on different proteins in the same network. For instance, proteins within a network can vary in the number and type of interactions, the position within the network (e.g., central versus peripheral), and the overall number of incorporated proteins in the network. Assessing the potential role of these factors, especially with multi-species comparisons, is crucial for understanding protein network evolution. Early comparative genomics studies, such as by Huynen *et al*. (Huynen et al. 1999), indicate that protein networks, even those involved in fundamental biological processes such as energy release by means of the citric-acid cycle, are characterized by variation in the composition and presence of specific proteins. Protein networks can change compositions by losing proteins or including novel proteins brought forward, for example through duplication events through neo- or subfunctionalization (Evlampiev & Isambert 2008; 2007). They can compensate for loss of proteins and new functions can evolve (Schüler & Bornberg-Bauer 2010). By inferring the evolutionary dynamics of protein networks, various patterns have emerged. Proteins with many interactions in the network (‘hub’ proteins) evolve slower than average (Fraser et al. 2002; Kim et al. 2006), especially when they use multiple binding interfaces (Kim et al. 2006). Interacting proteins evolve at similar rates (Fraser et al. 2002). Overall, protein networks are characterized by conservation in topology and function, but also by substantial levels of divergence in network constitution among species (Liang et al. 2006; Vleugel et al. 2012). Protein networks are thus affected by a combination of factors, including positive selection, selection on protein function or structure and drift (Pál et al. 2006).

In this paper, we examine a fundamental protein network with the aim to elucidate its evolution and to identify the factors that contribute to it. We focus on polarity establishment, a process essential for proliferation in basically all unicellular and multicellular organisms. Polarity establishment, or the asymmetrical distribution of cellular components, has been described in great detail in the budding yeast *Saccharomyces cerevisiae* (Chang & Peter 2003; Martin & Arkowitz 2014; Madhani 2007). To polarize, cells need to break up the symmetrical distribution of cellular content and self-organize in a polarized way. The small GTPase, Cdc42, is a central key protein in this process (Etienne-Manneville 2004; Park & Bi 2007; Johnson 1999). The asymmetrical distribution of so-called polarization proteins, recruited by Cdc42, determines the site of local growth, or budding in the case of the budding yeast *S. cerevisiae*, which is essential for proper cell division and mating.

Cdc42 is a highly conserved protein throughout eukaryotes at both the sequence and functional level (Johnson 1999; Martin 2015) and its activity is regulated through well-documented feedback mechanisms (Martin 2015; Goryachev & Pokhilko 2008; Irazoqui et al. 2003; Wedlich-Soldner et al. 2003). The proteins that directly interact with Cdc42 can be divided into five groups: the GTPase activating proteins (GAPs), that hydrolyze GTP to GDP and change Cdc42 to its inactive state; the guanine nucleotide exchange factors (GEFs), that catalyze the exchange of GDP for a new GTP molecule which activates Cdc42; the GDP dissociation inhibitors that extract Cdc42 from the membrane (Rdi1 is the only GDP dissociation inhibitor in budding yeast (Richman et al. 2004)); proteins involved in regulatory mechanisms, such as positive feedback (e.g., the scaffold protein Bem1 (Butty et al. 2002)); and a wide range of Cdc42 effector proteins which are activated by the active GTP bound state of Cdc42 (Figure 1A). Examples of Cdc42 effector proteins are the p21-associated kinases (PAK) Ste20, Cla4 and Skm1 (Johnson 1999), and the GTPase Interactive Components Gic1 and Gic2 (Brown et al. 1997). These proteins co-localize with Cdc42 during polarity establishment and form a protein complex by recruiting other proteins that are needed for proper actin and microtubule polarization (Johnson 1999; Brown et al. 1997; Drees et al. 2001). Various factors that could influence protein network evolution, such as the number of genetic and/or physical interactions and expression levels, have been determined for the proteins in this network in budding yeast (see Figure 1B). We therefore investigate the protein network evolution of polarization establishment among the ecologically and genetically highly diverse clade: the Fungi (Galagan et al. 2005; Ebersberger et al. 2012; Mueller & Schmit 2007).

**Fig. 1.**
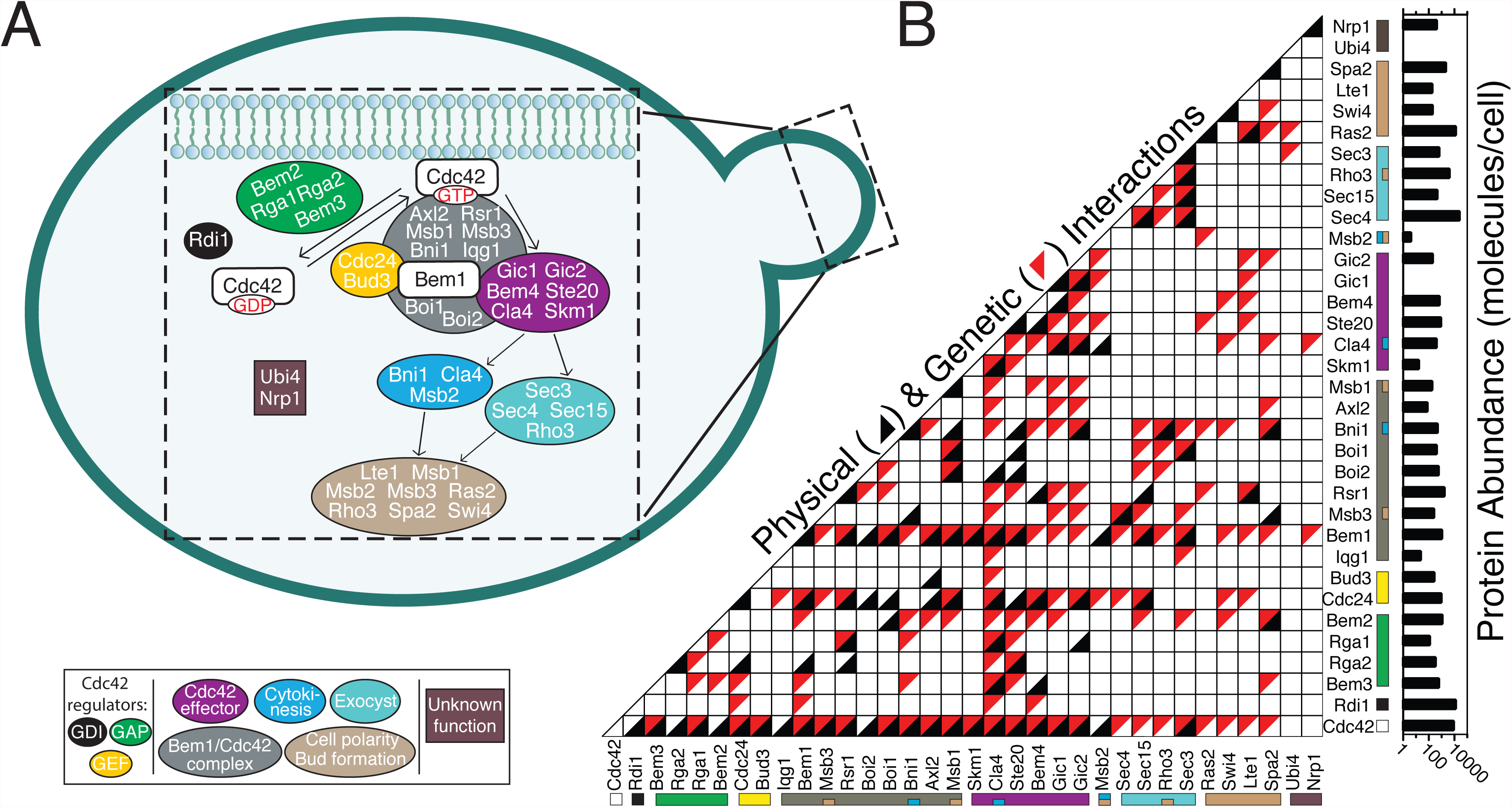
The central part of budding yeast’s polarization protein network. (A) Polarity establishment and subsequent budding takes place at one side of the budding yeast cell (cartoon). Insert: schematic overview of the 34 selected proteins and their functional groupings based on SGD; http://www.yeastgenome.org/, (Drees et al. 2001; Chang & Peter 2003; Martin & Arkowitz 2014; Madhani 2007). Cdc42 cycles between an active membrane-bound state (GTP) and an inactive cytosolic state (GDP). Depicted are the Cdc42 regulators and effectors, the Bem1/Cdc42 protein complex and several downstream steps. Nrp1 has a presumed function in polarity establishment (see Laan et al. 2015), Ubi4 has a described epistatic interaction with Cdc42 (BioGRID; thebiogrid.org). (B) Matrix of the genetic (in red) and physical (in black) interactions between the 34 selected polarization proteins based on SGD protein data. Proteins are color coded with the functional groupings from the (A) panel. Protein Abundance following Kulak et al. 2014 is displayed in the most right panel. Note that for Gic1 and Ubi4 no expression data was available.

The eukaryote kingdom of fungi is estimated >760 million years old (Lucking et al. 2009) and consists of up to 5.1 million estimated extant species (O’Brien et al. 2005). It includes an abundance of species with ecological, agricultural, medical and scientific relevance. Lifestyles can be restricted to a unicellular lifestyle, either yeast-like or non-yeast as observed in the basal clade of Microsporidia, or multicellular (i.e., filamentous species), or can consist of different stages, switching between two or more lifestyles (i.e., di-, trimorphic species). The wealth of different ecologies together with the available genomic and phenotypic resources and tools, such as the Saccharom*y*ces Genome Database (www.yeastgenome.org), make fungi an excellent tool for comparative studies between ecologically highly diverse, but also relatively closely related species. A vast increase of available fungal genomic datasets, especially fueled by initiatives such as the Fungal Genome Initiative (Rhind et al. 2011), the FungiDB (Stajich et al. 2011) and the 1K fungal genomes project (http://1000.fungalgenomes.org/home/; see also Sharma 2016), took place in the last five years prior to this study and provides the desirable scale of data to reach high phylogenetic resolution.

Although the processes of cell polarity and morphogenesis have been studied extensively in *S. cerevisiae* (Bi & Park 2012; Chant 1999; Pruyne & Bretscher 2000; Pruyne et al. 2004; Park & Bi 2007; Drees et al. 2001; Chang & Peter 2003; Martin & Arkowitz 2014; Madhani 2007), it is unknown to what extent the network’s topology is conserved across the fungal phylogeny, mainly because only a small number of divergent species has been examined, which are characterized by variation in both network composition and phenotypes (Diepeveen et al.). Due to its fundamental function in cell proliferation, the polarization protein network is hypothesized to be a conserved system (Chang & Peter 2003; Pruyne et al. 2004). In fact, several members of the polarization protein network, such as Cdc42, Cdc24 and Sec15, are found to be essential in *S. cerevisiae* (Liu et al. 2015). Interestingly, previously we showed that, under laboratory settings, the polarization network in *S. cerevisiae* is able to adapt to genetic perturbations to one of the core proteins: Bem1, which regulates Cdc42 (Laan et al. 2015). It is unknown to what extent this represents adaption under natural conditions. Thus, there is some information available on the conservation and evolvability (i.e., the ability of a species to evolve adaptive diversity) of a small number of individual polarization proteins, but a larger screen to quantify the evolutionary conservation across large phylogenetic distances is currently lacking.

Here we take advantage of the (newly) available tools and data in order to untangle patterns of protein network evolution with high phylogenetic resolution within a single, but phenotypically diverse kingdom. We aim to elucidate lineage-specific, independently recurrent and/or conserved patterns of protein network composition, and levels of protein conservation (i.e., both the presence of the protein in a species and sequence conservation) and divergence of 34 polarization loci among 200 closely related fungal species. We aim to elucidate factors involved in evolution of this particular and fundamental protein network.

## Material and Methods

### Focal (non-)polarization protein list and selected strains and species

We selected 34 polarization proteins (see Supplementary File 1, Figure 1) based on described physical or genetic interaction with the small GTPase, Cdc42, a key regulator of polarization (Etienne-Manneville 2004; Park & Bi 2007) and/or described functions of the protein in the polarization network on the Saccharomyces Genome Database (SGD; www.yeastgenome.org; June 2015; Cherry et al. 2011). For each of these proteins we downloaded the amino acid sequences from the SGD for *Saccharomyces cerevisiae* and used these sequences as reference or input query in the subsequent analyses (described below).

We obtained the full proteomes of 407 available fungal strains and species from Ensembl Fungi release 27 (June 2015; Supplementary File 1; Kersey et al. 2016) and created an initial proteome database (PDB) with the BLAST command line applications (version 2.2.31; Neumann et al. 2014). The proteome of *S. cerevisiae* (strain ATCC 204508/S288c) was downloaded from UniProt (www.uniprot.org). To perform the phylogenetic and reciprocal BLAST search analyses in an effective manner, we discarded strains and species with low quality proteomic data. We ran in-house scripts to determine the ‘optimum’ number of species that shared an ‘optimum’ number of communal proteins. In short, we calculated the number of shared protiens across the range of the 407 initial strains and species. We selected an ‘optimum’ of 200 strains and species (Supplementary File 1) for which we constructed the final PDB based on their proteomes as described above. The 200 strains and species had 86 proteins in common (Supplementary File 1). Note that these 86 proteins do not include any of the 34 polarization proteins. These protein sequences were used for the phylogenetic tree construction. We also constructed a *S. cerevisiae* (S288c) specific PDB for the reciprocal BLAST analysis.

### Phylogenetic tree construction

We retrieved the amino acid sequence of the 86 shared proteins (combined length of 49484 aa in *S. cerevisiae*) for the 200 strains and species. Alignments were made for each of the 86 proteins individually with Clustal Omega 1.2 (Sievers et al. 2011). The total length of the full alignment was 158856 aa. We performed phylogenetic inference on a single concatenated data file by means of the approximately maximum likelihood method as implemented in FastTree (version 2.1.8; Price et al. 2009) with the JTT model of amino acid evolution (Jones et al. 1992). The tree is mid-point rooted. The tree was visualized and edited in FigTree v1.4.2 (http://tree.bio.ed.ac.uk/software/figtree/). We checked the obtained support values and only reported them in the phylogeny, when they are < 0.9.

### Polarization protein conservation matrix

We screened the final PDB (see above) for the presence of the 34 focal polarization proteins through Reciprocal BLAST searches. We used the *S. cerevisiae* (strain ATCC 204508/S288c) amino acid sequence as input query. This was done to overcome computational limitations with orthology prediction between the 200 species. Other tools, such as DIAMOND (Buchfink et al. 2014) may ameliorate computational problems, but will result in a loss of sensitivity which is required when examining such specific protein networks. We performed local BLASTP searches without e-value constraints to be able to find hits, even in highly diverged species. We took the associated protein sequence of all obtained hits and performed local BLASTP searches again now against the *S. cerevisiae*-specific PDB, again without e-value constraints. We selected the hit with the best e-value in the initial BLAST search and called the hit a ‘true match’ only if the same protein ID was retrieved as the original *S. cerevisiae* query. If there was no match in protein ID between hit and query, then we called it a ‘no match’. For the ‘true matches’ we corrected the similarity scores (i.e., the number of positive-scoring matches) of the best BLAST hit to the query protein length of *S. cerevisiae*, thereby obtaining the fraction of similarity per hit. For the ‘no-matches’ we assigned a similarity score of 0. We generated a matrix of similarity scores by combining all the obtained scores and organizing them according to the species order as observed in our constructed phylogeny for each protein. These steps were performed with in-house python scripts. The generated data matrix was displayed as gray-scale matrix in R version 3.1.2 (R Core Team 2014).

We constructed a separate protein matrix for the group of 36 strains/species with high genome quality (i.e., for which the number of chromosomes is known and the number of scaffolds (roughly) equals the number of chromosomes; Supplementary File 1). For these species high quality genomes are available and therefore, overall genome quality should not affect our obtained results by means of incomplete genome sequences, and thus the possibility of absent protein sequences.

To determine if there was a group of proteins systematically present in a high number of species, we plotted the overall prevalence (%) across the 200 strains/species for each protein. We also plotted the overall prevalence of the two main fungal lineages Basidiomycota and Ascomycota separately in order to make sure that the proteins were present in high prevalence in both major clades. We used a 70% cut-off value in both clades as criteria for high prevalence and call the proteins that meet these criteria the conserved core of polarization proteins. We based this cut-off value on the observation that there is a gap in prevalence between 45% and 70% for the full dataset of 200 strains/species, dividing proteins into two groups. We also plotted the difference in prevalence between the Ascomycota and the Basidiomycota, to depict proteins that are particularly prevalent in either group.

### Statistical analyses

We tested for a potential correlation between our obtained pattern of matches (i.e., the total number of ‘true matches’ per strain/species) and genome quality for which we used two assembly statistics. We obtained the number of contigs and N50 of contigs (i.e., length of contigs constituting 50% of the bases in the assembly), the number of scaffolds and the associated N50 for each of the genomes of the 200 selected strains/species from the European Nucleotide Archive (http://www.ebi.ac.uk/ena; Leinonen et al. 2010). Data was tested for normality with D’Agostino & Pearson omnibus normality tests as implemented in GraphPad Prism version 5.0 for Mac OS X (GraphPad Software, La Jolla California USA, www.graphpad.com). Correlations were tested with a spearman’s rank correlations as implemented in GraphPad Prism.

We also tested if genome quality affected our obtained results of the full protein matrix by directly comparing the observed number of proteins per species clade-wise for four major lineages to the ones of the same lineages in the reduced matrix of 36 species. We used the Microsporidia, Saccharomycotina and Pezizomycotina lineages because these clades are also represented by at least 4 individuals in the reduced matrix. We divided the Saccharomycotina clade into two subclades because we observed distinctive lineage-specific patterns, such as gains/losses and distinctions in similarity scores, dividing them in two clades. Saccharomycotina clade 1 represents the Saccharomycetaceae family and the Saccharomycotina clade 2 includes the Debaryomycetaceae, Pichiaceae and Phaffomycetaceae families. Data was tested for normality with D’Agostino & Pearson omnibus normality tests as implemented in GraphPad Prism. We performed a Kruskal-Wallis test comparing the medians of the two matrices per defined lineage.

To test hypotheses about the cause(s) of differences observed in prevalence between the core and non-core proteins, we tested if these groups of proteins differed in number of genetic interactions, number of physical interactions and overall abundance of the proteins, as indicative for gene expression level. Data for the number of interactions of the 34 proteins with other members in the network in *S. cerevisiae* were obtained from SGD. We gathered information about protein abundance (molecules/cells) in *S. cerevisiae* from Kulak *et al.* (Kulak et al. 2014). Data was tested for normality with D’Agostino & Pearson omnibus normality tests as implemented in GraphPad Prism. We also included the prevalence in the analyses. We performed, a Kruskal-Wallis test (for interactions) and a Mann Whitney tests (for prevalence and protein abundance) with GraphPad Prism.

### Multiple Factor Analysis

Because we expected multiple continuous and categorical variables to potentially co-vary and correlate with the observed number of proteins, we performed Multiple Factor Analysis (MFA) on a dataset of the 200 strains/species. We included the following variables: proteins (i.e., the total numbers of proteins as observed in the full protein matrix), genome quality (i.e., the number of Contigs and N50 of Contigs), Lineage (i.e., the main retrieved phylogenetic clades: Microsporidia, Cryptomycota, Wallemiomycetes, Pucciniomycotina, Ustilaginomycotina, Agaricomycotina, Taphrinomycotina, Saccharomycotina (clades 1 and 2, as defined above), Pezizomycotina), genetic distance (in respect to the reference *S. cerevisiae*), and lifestyle (i.e., unicellular, yeast, filamentous, dimorphic yeast-filamentous, dimorphic yeast-pseudohyphal, and trimorphic). We calculated the genetic distance between the examined strains/species and the reference. We used the concatenated amino acid sequences of the 86 proteins from the phylogenetic analyses (158854 aa; see above) and calculated the genetic distance by using the JTT model of amino acid evolution in MEGA 7: Molecular Evolutionary Genetics Analysis version 7.0 for bigger datasets (Kumar et al. 2016). We obtained the lifestyle information from the Fungal Databases of the CBS-KNAW Fungal Biodiversity Centre (http://www.cbs.knaw.nl/) and literature (Nagy et al. 2014; Bastidas & Heitman 2009; Gauthier 2015). We performed the MFA with the FactoMineR R package version 1.33 (Lê et al. 2008) package in R version 3.3.2 (R Core Team 2014) under Rcmdr version 2.3-2 (Fox & Bouchet-Valat; Fox 2005; 2016). We used 3 quantitative groups: genetic distance, genome quality (i.e., 2 variables) and proteins, and two qualitative groups; lifestyle and lineage. Continuous variables were scaled and standard settings were used. We first checked the eigenvalues for the first ten dimensions to determine the appropriate number of dimensions to consider. In particular we checked for a drop in decline in variance (i.e., broken stick method; Jackson 1993). Length and directions of continuous variables were plotted onto the first two dimensions and were visually checked. Partial axes for the fist first three dimensions were visually checked. The five groups were plotted onto the first two dimensions. We plotted individuals onto the first two dimensions and color-coded them according to lineage.

## Results

### Phylogeny of 200 fungal strains/species is highly supported

In order to examine protein network evolution of fungal polarity establishment, we first estimated the phylogenetic relationship for our focal species. We inferred the phylogeny by means of the approximately maximum likelihood method on 86 homologous non-polarization proteins that these 200 strains/species have in common (see methods for details; total alignment length: 158,856 amino acids (aa; Figure 2)). Our phylogenetic tree is well-resolved and highly-supported and includes four main monophyletic phyla: the club fungi and relatives (Basidiomycota); sac fungi (Ascomycota); and the two basal phyla Cryptomycota and Microsporidia. Within the Basidiomycota, we found 100% support for the monophyletic subphyla Ustilaginomycotina, Pucciniomycotina, Agaricomycotina and Wallemiomycetes. Our observation of a basal position of the Wallemiomycetes species is in accordance with previous findings (Matheny et al. 2006; Zalar et al. 2005; Hibbett et al. 2007), although its exact position within the Basidiomycota phylum, and in relation to the Agaricomycotina in specific, has been varying (Matheny et al. 2006; Padamsee et al. 2012). Within the Ascomycota, we found full support for the monophyly of the Taphrinomycotina, Saccharomycotina and the Pezizomycotina, consistent with previous findings (Hibbett et al. 2007; Schoch et al. 2009). A discussion on relationships of deeper braches and clades is, however, beyond the scope of this work, although we observed almost exclusively high support values (i.e., >0.9) for the deeper branches and many relationships agree with previously published work (e.g., Saccharomycotina lineage; Shen et al. 2016).

**Fig. 2.**
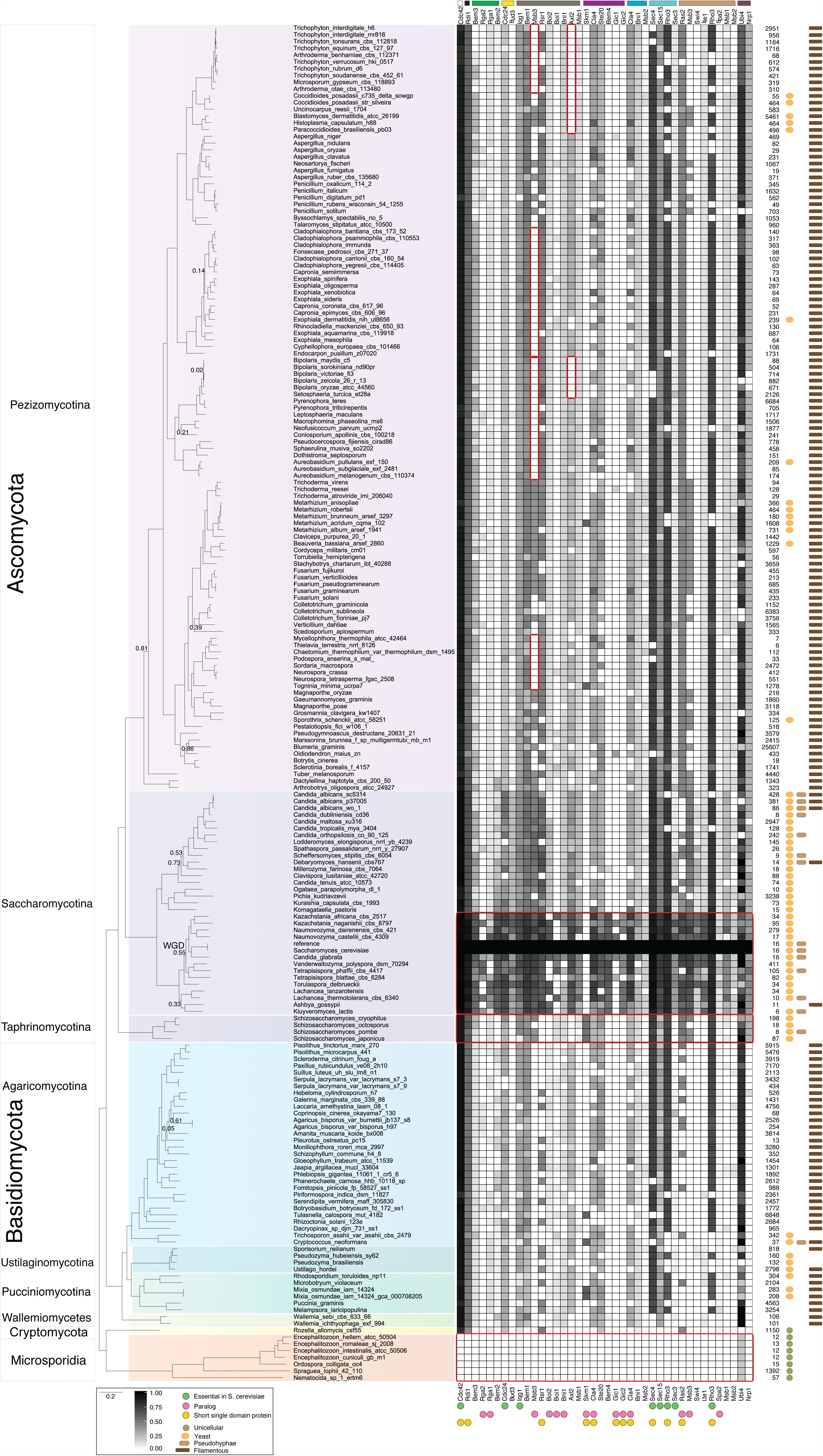
Phylogenetic relationships between the 200 fungal strains/species and the protein matrix for the 34 selected polarization proteins. The phylogeny is based on 86 protein sequences (158856 aa) and we used the approximately maximum likelihood method and the JTT model of amino acid evolution. Support values are almost exclusively above 0.9, except when shown on the tree (12 instances). The tree includes the phyla: Microsporidia (in orange), Cryptomycota (in yellow), Basidiomycota (in blue) and the Ascomycota (in purple). Subphyla are shades of the same phylum color. Phylogenetic relationships greatly follow known relationships (Matheny et al. 2006; Zalar et al. 2005; Hibbett et al. 2007; Schoch et al. 2009; Shen et al. 2016). The protein matrix displays the similarity scores of the reciprocal BLAST search. White fields represent no match of the query protein in the respective PDB; black field represent a match with 100% similarity score; grey fields represent a match with <100% similarity score. Proteins are ordered and color coded following Figure 1A. Essential proteins (in *S. cerevisiae*), paralogs and short single domain proteins are labeled with green, pink and yellow bullets at the bottom of the matrix. Various recurrent and lineage-specific patterns, as discussed in the main text, are highlighted by red outlines. Genome quality as in number of contigs is shown in the most right column, followed by the life styles of the fungal species (cartoons: yeast-like (orange), non-yeast-like unicellular (green), pseudohyphal (light brown), filamentous (dark brown)).

### Genome incompleteness has little impact on the protein matrix

We constructed a protein matrix consisting of the 34 polarization proteins and 200 strains/species based on our reciprocal BLAST approach (see methods for details). This way we can both determine the presence of a protein in the examined strains/species and, when present, the level of sequence similarity in respect to *S. cerevisiae*.

Our approach resulted in a detailed protein matrix indicating the presence, level of divergence in respect to *S. cerevisiae*, and absence of the 34 polarization proteins in the examined strains/species (Figure 2). To determine if the quality of the genomic resources had an effect on the obtained results, we tested if there was a correlation between the total number of contigs, the N50 of contigs (i.e., length of contigs constituting 50% of the bases in the assembly) and the total number of proteins we obtained per strain/species. We observed that the quality of the 200 selected strains/species’ genomic resources was highly variable (Supplementary File 1). Contig N50 range from 6 kb - 9 Mb. Genomic resources with short N50 may suffer from missing data such as missing exons and/or gaps (English et al. 2012), which could include, or result in, missing loci. A recent survey of >200 fungal genomes indicates that potentially only 40% reach the set cut-off for representative completeness (Cisse & Stajich). We found weak but significant correlations between the total number of obtained proteins per species and the number of contigs (Spearman rho = -0.29, P-value < 1.0 x 10^−4^), and the contig N50 (R = 0.32, P-value < 1.0 x 10^−4^; Figure 3). Thus, we potentially have cases of false negatives, or mismatches, in our protein matrix. In order to examine this pattern in more detail and to diminish the potential effect of missing proteins on our reciprocal BLAST analysis, we selected 36 strains/species with the most complete genomes (i.e., the number of scaffolds roughly equals the number of chromosomes) and constructed a reduced matrix (Figure 4). We then compared the overall number of observed proteins in the full matrix with the reduced matrix for all members of four lineages: Microsporidia, two Saccharomycotina clades, and the Pezizomycotina (Supplementary File 2). We did not observe significant differences in the number of proteins between the complete matrix and the reduced matrix (P-value > 0.05 for Microsporidia, Saccharomycotina I, Saccharomycotina II, Pezizomycotina), indicating that the potential effect of missing data in our full matrix is, if any, minimal. Next, we examine the reduced matrix for patterns of protein conservation and divergence and compare them to the original matrix in order to test their generality.

**Fig. 3.**
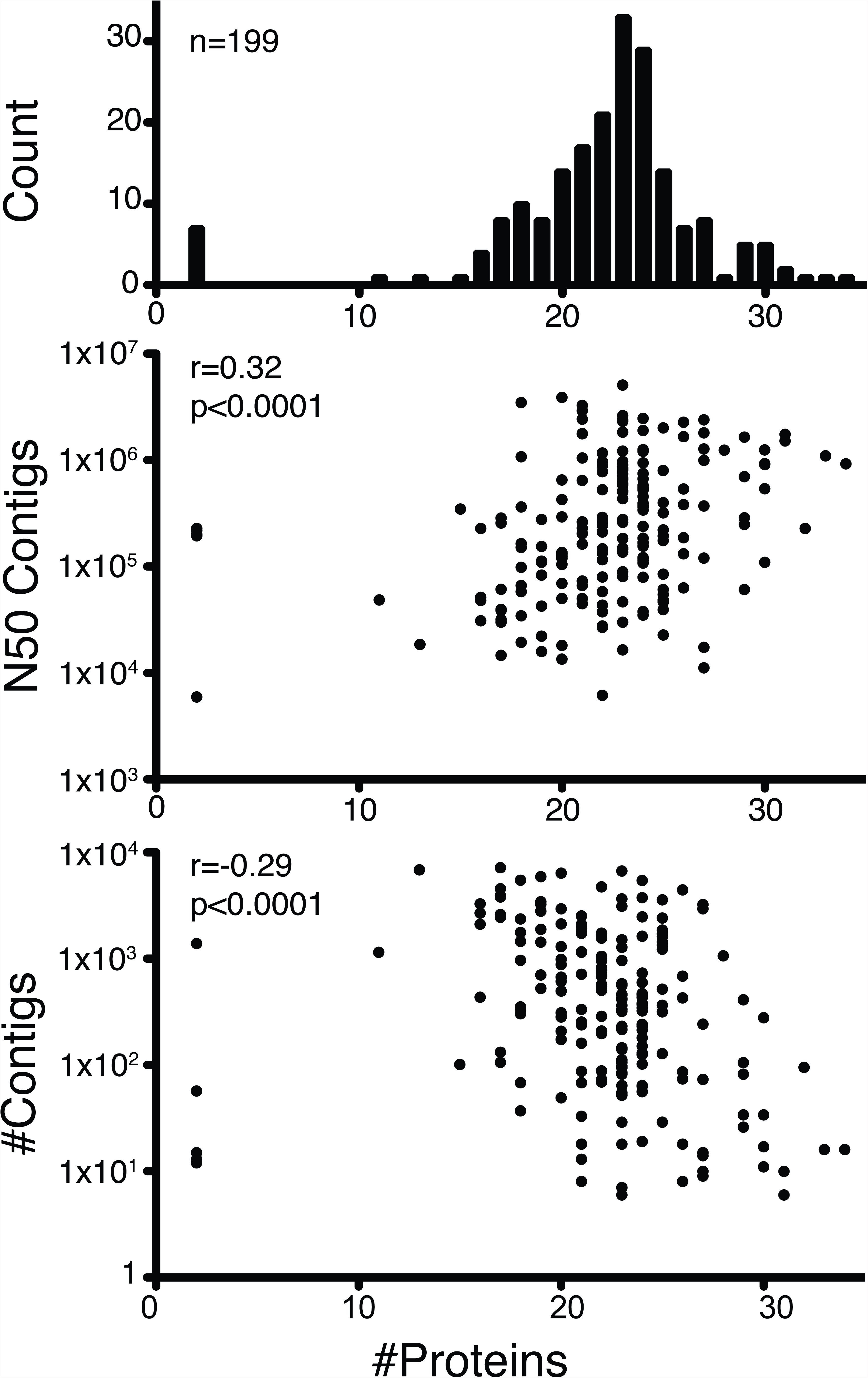
Correlation between genome quality and number of retrieved proteins. The top panel (Count) shows the distribution of strains/species for the number of retrieved proteins. The center panel shows a statistically significant positive correlation between the N50 of contigs of the genome and the number of retrieved proteins. One data point (x=11, y=9.5x10^6^) was omitted from the plot for clarity of the plot. The bottom panel shows statistically significant negative correlation between the number of contigs in the genome and the number of retrieved proteins. One data point (x=12, y=25607) was omitted from the plot for clarity of the plot.

**Fig. 4.**
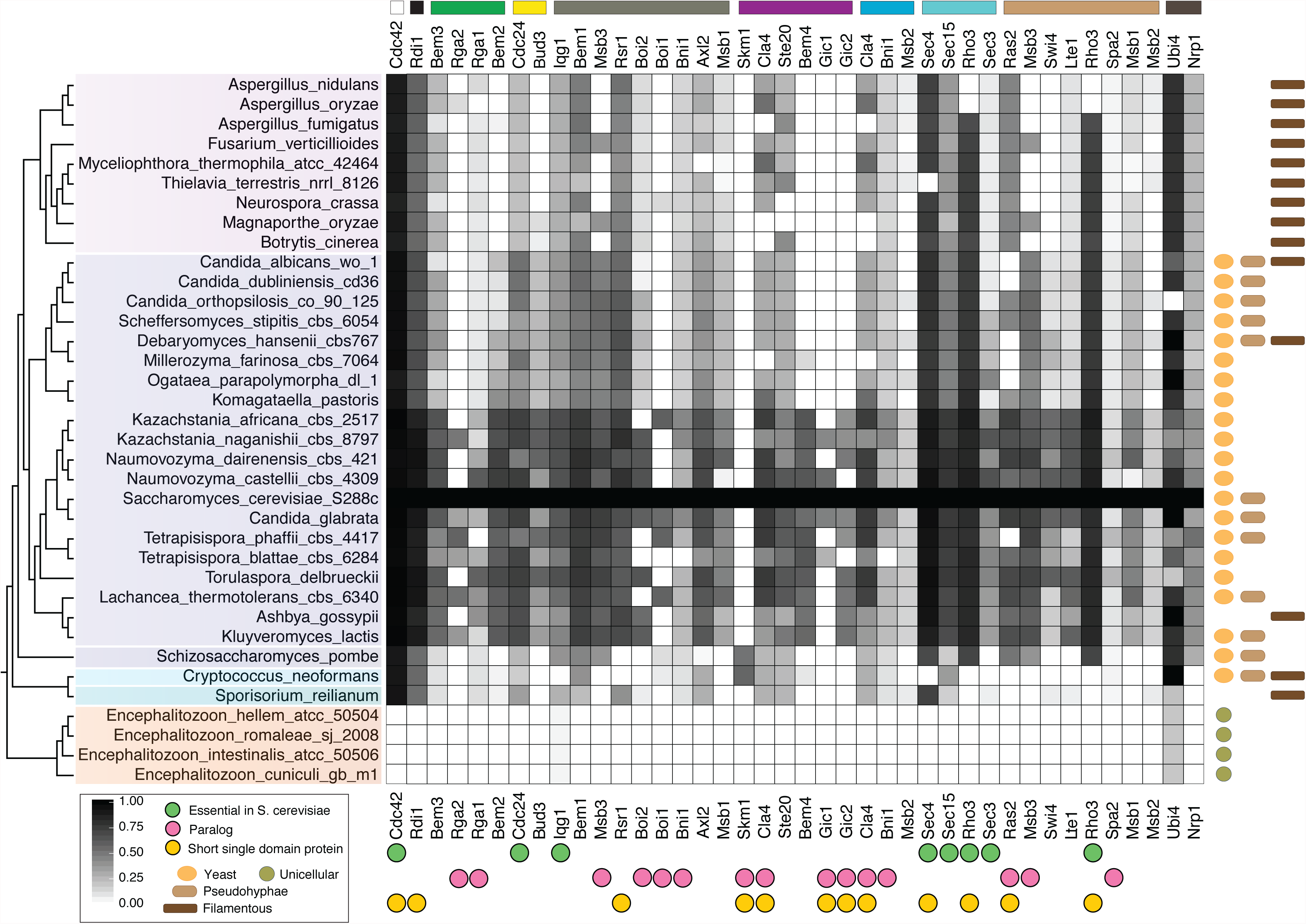
Protein matrix of the 36 species with highest genome quality. The matrix displays the similarity scores of the reciprocal BLAST search for species of the Microsporidia (in orange), Basidiomycota (in blue) and the Ascomycota (in purple). The cladogram on the left represents the phylogenetic relationship between strains/strains based on Figure 2 (note: the branch lengths are fixed and do not represent amino acid substitutions). Proteins are ordered and color coded following Figure 1A. Essential proteins (in *S. cerevisiae*), paralogs and short single domain proteins are labeled with green, pink and yellow bullets at the bottom of the matrix. The life styles of the species are depicted on the right column.

### The polarization protein network is dynamic and has a conserved core

To examine the fungal polarization network for patterns of protein network evolution, we screened the protein matrices for variation in protein prevalence (i.e., the overall number of species a protein is present in), levels of protein divergence and the repertoire of proteins (i.e., the composition of polarization network per species). We observed variation in the polarization network at different levels and magnitudes. We observed great variation at the level of amino acid similarity between different proteins, across different strains/species and between different lineages (Figure 4, Figure 2). For instance, Cdc42 is present with high levels of similarity in all species except in the Microsporidia, where it is not present in any of the examined species. Cla4, Ste20 and Cdc24 are found throughout the phylogeny as well, but their similarity scores vary greatly across species. These patterns were observed both in full and reduced matrices.

Next we examined the variability of the composition of the polarization protein repertoire between strains/species. We assessed the overall number of different protein combinations and the total number of unique repertoires across the reduced matrix (36 strains/species) and the full matrix (200 strains/species). We observed nearly identical fractions for the total number of different protein repertoires for the two matrices (i.e., 28/36 and 155/200). This indicates that the composition of repertoire is highly variable, with many different observed combinations of proteins. For both matrices the overall fraction of unique repertoires (i.e., protein repertoires observed in a single species) was also very similar, 0.64 (reduced matrix) and 0.67 (full matrix). We thus find that the majority of species are characterized by a unique set of polarization proteins not found in other species. Interestingly, we observed several specific combinations multiple times. For instance, we observed the same pattern for all 7 Microsporidia species (Iqg1 and Ubi4). We observed most cases of repeated repertoires in the species-rich and well-covered lineages Pezizomycotina (113 species in the full matrix) and Saccharomycotina lineages (20 species in the reduced matrix) (Supplementary File 3). These combinations include prevalent and functionally diverse proteins such as Rdi1, Bem1/2/3, Bni1, Axl2, Cla4, Ste20, Sec3/4/15, and Msb1/3.

To examine the overall prevalence of each protein across the 200 strains/species in more detail, we screened the full matrix (Figure 5). We observed 23 proteins that were present in ≥ 70% of all examined species (e.g., Iqg1), seven proteins are more commonly found in the Basidiomycota (e.g., Bem2), and four proteins restricted to the Ascomycota (e.g., the paralogs Msb3, Gic1 and Gic2). Most proteins are highest prevalent in the Ascomycota (Figure 5 top panel). We observed a perceived threshold at ~60% prevalence for proteins across all species examined that clearly divided the dataset (Figure 5). We found 11 proteins that are present in less than 45% of the 200 strains/species, while the other 23 proteins are present in at least 70%. We used this 70% mark as cut-off value to determine conserved proteins (i.e., based on prevalence) in both Ascomycota and Basidiomycota, individually. Using this, we excluded those proteins that are more or less Ascomycota-specific, such as Nrp1. We observed the following 16 proteins in high prevalence in both Ascomycota and Basidiomycota: Cla4, Axl2, Rho3, Boi2, Ste20, Sec4, Bni1, Bem3, Spa2, Cdc24, Sec15, Ubi4, Bem1, Cdc42, Rdi1 and Iqg1. We called these proteins the conserved core of polarization across fungi. We found this full repertoire of core proteins in 66 out of 200 strains/species and >95% of all strains/species had a protein network consisting of 12 or more core proteins (Supplementary File 3).

**Fig. 5.**
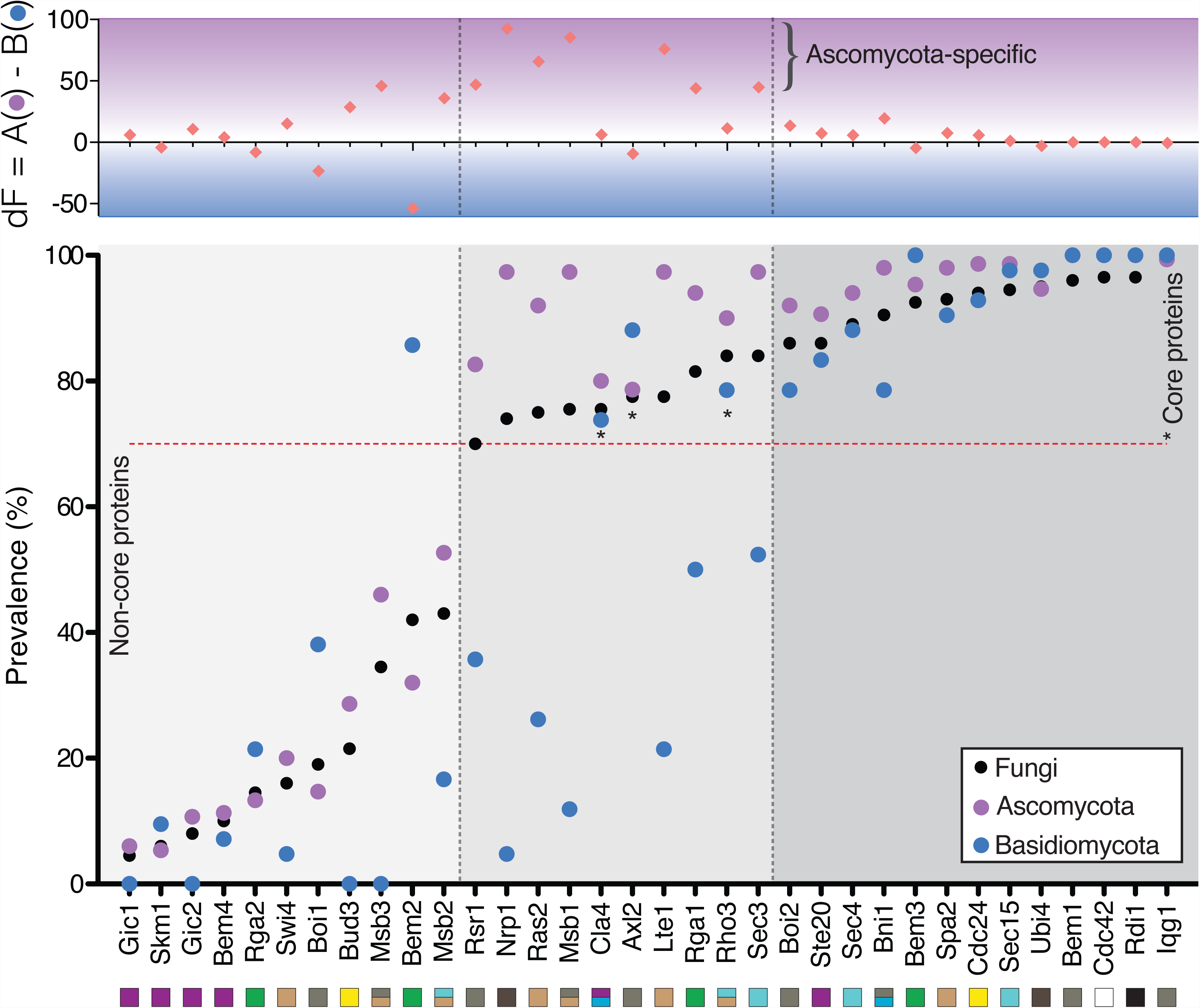
Polarization proteins prevalence. Prevalence of the 34 polarization proteins for all examined fungal species (black circles), the Basidiomycota species (blue circles) and the Ascomycota species (purple circles). Proteins are ordered based on their overall prevalence in all examined strains/species. The 70% criterion is marked by a horizontal red dotted line. Shading in the bottom part reflects grouping of proteins with < 70% prevalence in all strains/species (light grey; left), proteins with prevalence > 70%, but prevalence in the Basidiomycota is not in all cases >70% (grey; center), proteins with >70% prevalence in all examined groups (i.e., core proteins; dark grey; right). Core proteins in the middle group are marked by an asterisk (*). Difference in prevalence between the Ascomycota and Basidiomycota is presented in the top panel (pink diamonds).

### Core proteins have higher protein abundance but not more interactions

As we observed a group of proteins at high prevalence across clades, we tested if there is a correlation between this conserved core of proteins and factors known to influence protein (network) evolution, such as number of protein-protein interactions and expression levels. We thus tested the following hypothesis: core proteins are conserved because they are either functionally important and/or because they are present in high quantity.

Core proteins had higher prevalence than non-core proteins (Figure 6A). We found no significant difference in the number of either genetic or physical interactions (based on observations in *S. cerevisiae*) between the core proteins and the non-core proteins (Figure 6B). We did find a barely significant difference in protein abundance (as measured as molecules per cell in *S. cerevisiae*; Kulak et al. 2014) between the core proteins and non-core proteins (p=0.03; Figure 6B). Core proteins have higher protein abundance than non-core proteins.

**Fig. 6.**
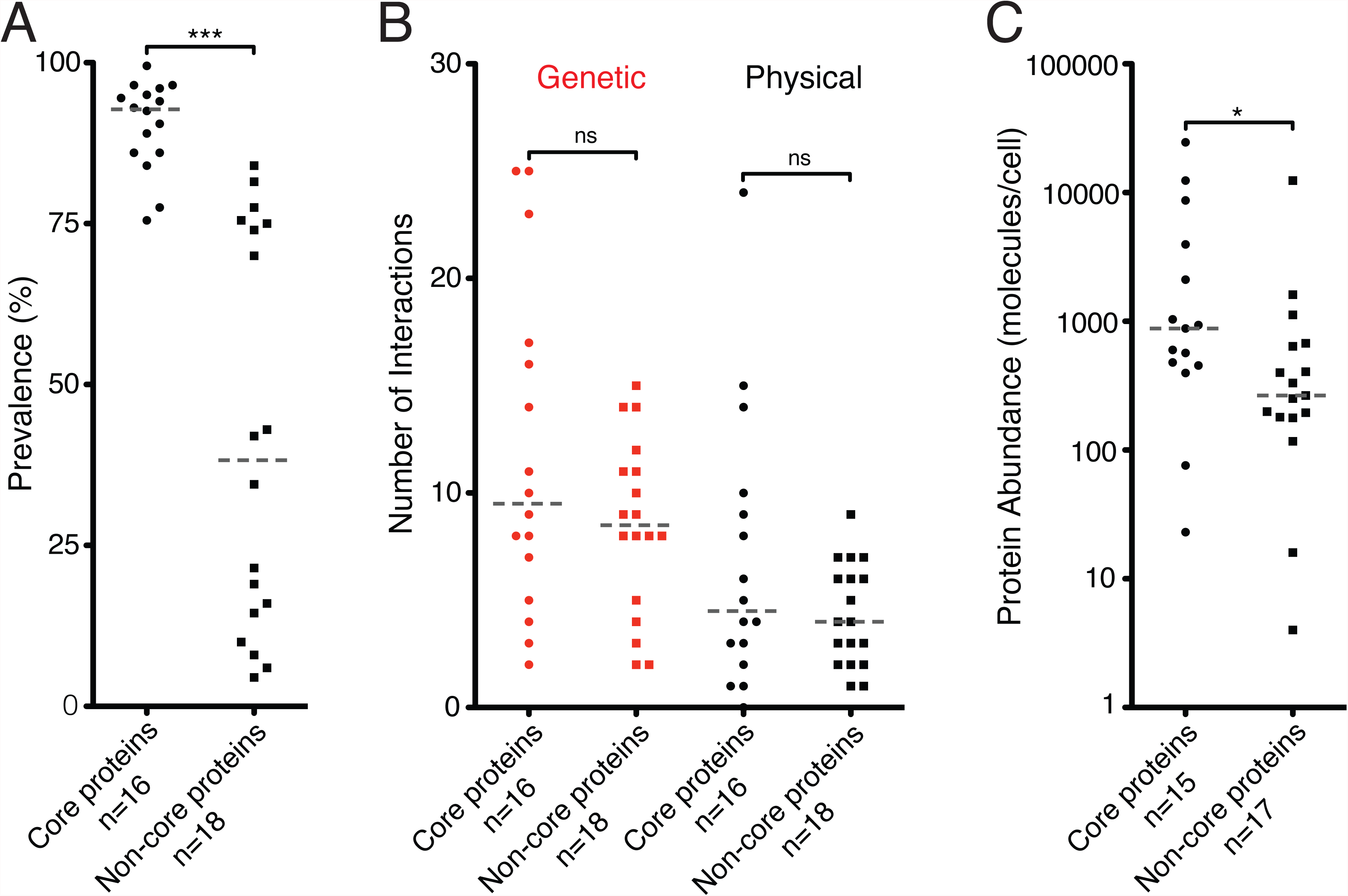
Comparison between the core and non-core proteins. (A) Significant difference in the observed prevalence of the core and non-core proteins (P-value < 0.0001). (B) Number of genetic interactions (in red) and physical interactions (in black) between the 34 examined polarization proteins. No difference was observed between the core and non-core proteins in the number of genetic or physical interactions. Data was obtained from SGD protein data. (C) Significant difference in protein abundance between the two groups. Core proteins have higher protein abundance (P-value = 0.03). Note that data for Gic1 (non-core) and Ubi4 (core) was unavailable. Core proteins are depicted as circles, while non-core proteins are depicted as squares. Grey lines depict medians.

### Lineage, lifestyle and genetic distance co-vary with protein network size

In order to characterize the 200 strains/species by different factors that could influence protein network evolution and in specific could correlate with the observed patterns of differences in overall number of polarization proteins, we performed Multiple Factor Analysis. We considered the following factors: size of protein repertoire (i.e., the total number of proteins we detected with the Reciprocal BLAST search per strain/species), lifestyle, lineage, genome quality (i.e., the number of contigs and the N50) and genetic distance to the reference *S. cerevisiae*, based on 86 shared proteins.

To determine the adequate number of dimensions to screen, we used the broken stick method (Jackson 1993). We found a drop in variance after the third dimension (Supplementary File 4), therefore we only considered the first three dimensions. Dimension 1 is constructed based on four groups: lineage (contribution is 27.44%), lifestyle (24.72%), genetic distance (23.39%) and proteins (23.23%). Dimension 2 and 3 are both based on lineage (44.16%; 50.61%) and lifestyle (43.87%; 46.31%). Together these three dimensions accounted for 40.70% of the variance in the data. Dimension 1 explained 20.28%, dimension 2 12.37% and dimension 3 8.06% (Figure 7D). We did not find a substantial (>0.5) contribution of genome quality, indicating that the number of contigs and/or N50 of contigs did not explain the variation in the protein repertoire and other examined factors. Supplementary File 4B shows that lifestyle and lineage vary together and that they vary together with protein repertoire and genetic distance. Supplementary Files 4C & 4D indicate that protein repertoire and genetic distance only correlate with dimension 1, while lineage and lifestyle also correlate with dimensions 2 and 3.

**Fig. 7.**
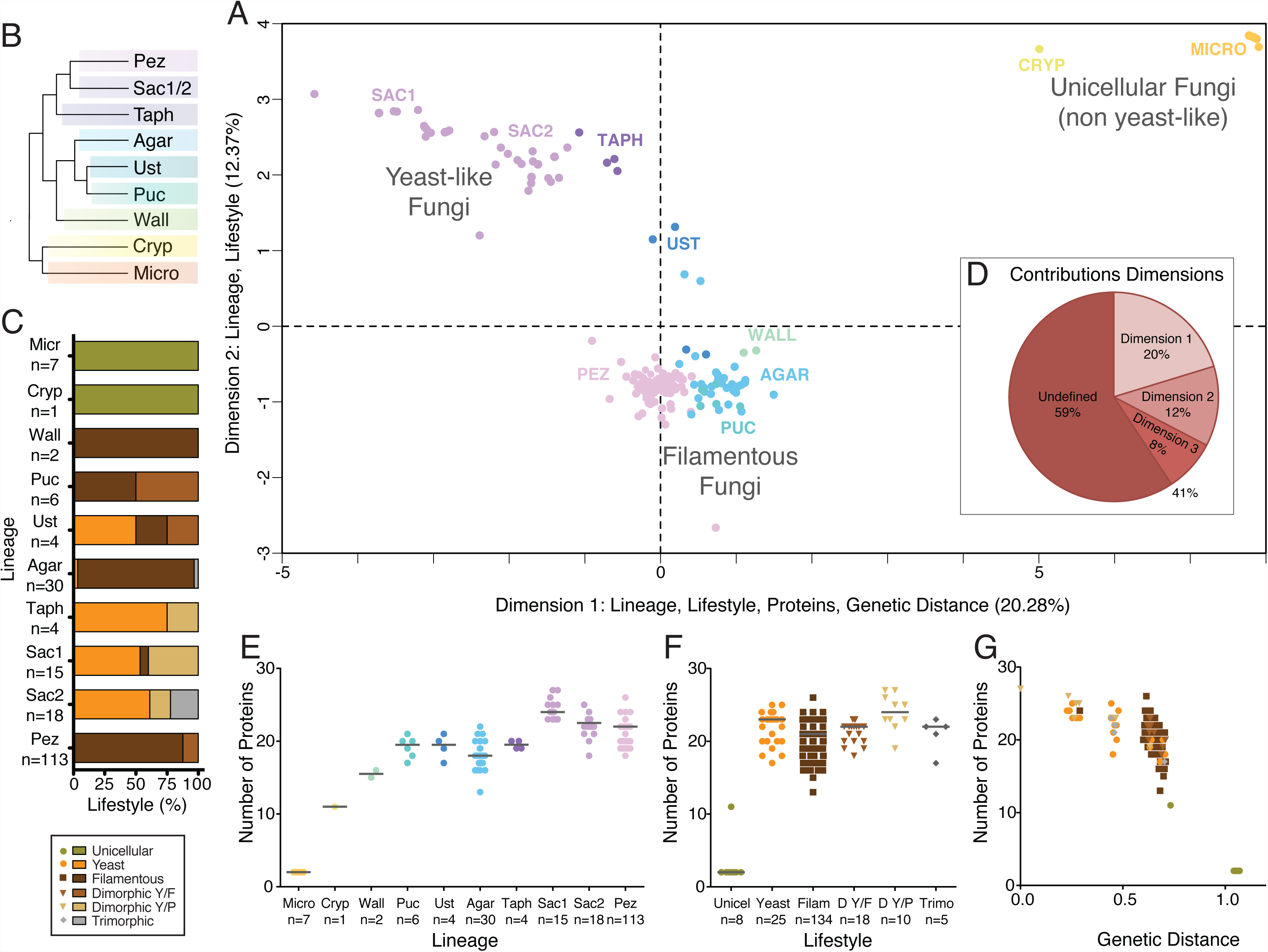
Multiple Factor Analysis and correlations. (A) Multiple Factor Analysis of the number of polarization proteins, lineage, lifestyle, genomic quality and genetic distance. The 200 strains/species are plotted and color-coded according to their phylogenetic lineage as in Figure 2. Dimension 1 explains 20.28% of the observed variation and the following four factors constitute to its construction (in order of importance): lineage, lifestyle, genetic distance, number of observed proteins. Dimension 2 explains 12.37% of the variation in the data and is based on the variables lineage and lifestyle. Main areas occupied by specific lineages are labeled accordingly for clarity. A clear distinction can be made between yeast-like fungi (left top corner), filamentous fungi (center lower part) and unicellular non-yeast like fungi (right top corner). (B) Cartoon depicting the topology of the major clades. Note that the two Saccharomycotina lineages are shown together. The length of branches do not represent observed branch lengths. See Figure 2 for full phylogeny. (C) The distribution of lifestyles (in percentages) for the ten different phylogenetic lineages. The number of strains/species per lineages is given. Lifestyles are color-coded as in legend at the bottom right of the figure. The 200 strains/species are classified as unicellular, yeast, filamentous, dimorphic (either yeast/filamentous or yeast/pseudohyphal) and trimorphic following Figure 2. (D) Pie plot depicting the percentage of variation explained by the three main dimensions. The three dimensions account for 41% of the observed variation, leaving 59% undefined. (E) The number of observed proteins in ten different phylogenetic lineages. Groups are color-coded per lineage as in Figure 2. Medians are given as grey lines. (F) The number observed proteins in the different lifestyles. Grey lines represent medians. (G) The number of observed proteins plotted versus the genetic distance (in respect to *S. cerevisiae*). Strains/species are color-coded according to their lifestyle morphology.

We plotted the 200 strains/species onto the first two dimensions to visually examine if they cluster to specific patterns based on e.g., morphology or descent (Figure 7A). As four factors mainly contributed to the axes, various patterns were observed. Overall, prevalence (proteins) declined horizontally (from left to right), with an increase in genetic distance. Lineage and lifestyle contributed to both axes, and a clear distinction could be made between three main clouds of individuals according to their morphology: the top left corner represents the yeast-like fungi, the top right corner represents the unicellular non-yeast-like fungi, and the lower center represents the filamentous fungi. The yeast-like fungi mainly constituted out of the yeast and dimorphic (Y/P) lifestyles, while the lineages that mostly consist of the filamentous lifestyle were found in the bottom center cloud (Figure 7C). We observed a split in the Ustilaginomycotina species, as two species are yeast-like and two species are filamentous. Interestingly, the Ascomycotan and mostly filamentous Pezizomycotina clustered together with the Basidiomycotan individuals in the filamentous group. This observation is not in line with the phylogenetic relationships between these clades, but it does indicate that variation in the factor lifestyle is more similar for the Pezizomycotina and the Basidiomycotan species, as opposed to the Pezizomycotina and other Ascomycotan species. We observed similar patterns when we examined these factors individually versus the protein repertoire (Figure 7E-G). Our results suggest that the Basidiomycota and the filamentous Ascomycotan species have a smaller repertoire than the yeast-like Ascomycota, and that the two basal lineages have an even smaller repertoire (Figure 7E). Lastly, we did observe a pattern of decreasing number of polarization proteins with greater genetic distance to our reference *S. cerevisiae* (Figure 7G).

## Discussion

Here we assess evolution of the fungal polarization network at high phylogenetic resolution to examine patterns of lineage-specific, independently recurrent and/or conserved patterns of protein network composition along with levels of protein divergence. We observe that the fungal polarization protein network is characterized by both strong protein conservation *and* variation in protein prevalence and sequence similarity. Our results indicate that while certain proteins are nearly always needed potentially for specific functions (i.e., functional conservation), various other functional steps seem to be fulfilled by a variable repertoire, indicating flexibility in the network composition. Below, we discus these observations in context of protein network dynamics, functionality of the protein network, and potential causal factors of protein network evolution.

### The fungal polarization network is highly variable at lineage-specific levels

It is clear that protein network evolution has a variety of outcomes, such as network expansion/reductions, interaction effects and protein divergence (Schüler & Bornberg-Bauer 2010; Voordeckers et al. 2015), brought forward by e.g., gen(om)e duplication, selection on protein function or structure and drift (Pál et al. 2006). We found that most proteins of the polarization network have high levels of divergence in amino acid sequence across fungi and that the specific build up of the protein repertoire per strains/species is highly variable. We find both variation at large phylogenetic distances, such as between subphyla, and between strains/species of the same clade. This indicates that, although the polarization network is involved in fundamental cellular functions across organisms, the network, as we know that in *S. cerevisiae*, is not a conserved entity. Work based on the first available fungal genomes reveal remarkable levels of divergence (Galagan et al. 2005), with even less than 50% similarity in amino acid sequence in comparisons of Ascomycotan species (Dean et al. 2005). Screening these genomes for networks reveal that especially regulatory pathways are recurrently characterized by substantial levels of variation, in that elements can be gained or lost over time (Tanay et al. 2005; Muñoz et al. 2016; Tuch et al. 2008; Habib et al. 2012). Our work provides further support for the eminent finding that proteomes and networks constantly change (Coulombe-Huntington & Xia 2017), not only in Ascomycota as previously shown, but also in Basidiomycota and basal lineages as Cryptomycota and Microsporidia.

The substantial levels of variation that we observed in the polarization network could be caused by the remarkable differences in how fungal species polarize and grow (e.g., isotropic, (a)symmetric). In fact, we do find a clear clustering of yeast-like fungi, non-yeast like unicellular fungi and filamentous fungi in our MFA plot. While budding yeast polarizes in a switch-like way, filamentous species are characterized by continuous hyphal growth and thus need a constant state of polarity. Differences at the protein levels between species with differences in polarization/growth mode have also been described. The Rho GTPase Rac1 has partly overlapping functions with Cdc42 in regulating polarization in a variety of filamentous species (Banuett et al. 2008), but not in *S. cerevisiae*. We believe that the observed levels of variation have a strong functional component.

To what extent does this high variability of the protein network affect functionality? As functional studies are not available for the majority of examined species, we made use of the functional classification of proteins of *S. cerevisiae* (see Figure 1). We found that over 90% of examined strains/species have at least one protein present from all nine defined functional groups. This could imply that the overall functional pathway of polarity establishment, by means of the regulation of Cdc42, is similar across the fungal tree. Exceptions to this overall pattern of consensus are observed throughout the tree (e.g., Microsporidia, *Penicillium rubens*, *Suillus luteus*, *Serpula lacrymans*, lacking e.g., Cdc42 effectors and/or GEF proteins). Further functional exploration of protein networks in additional non-model species is needed to determine the level of orthology of these networks.

### Variation in polarization network; from stark reductions to lineage-specific additions

We found high levels of lineage-specific patterns, of which various patterns coinciding with monophyletic clades. For instance, Msb3 and Axl2 are individually lost in various Pezizomycotina clades (Figure 2). The protein matrix also showed very similar patterns for the four Taphrinomycotina species, which are quite dissimilar from the other Ascomycota clades. These species have very distinct ecologies, as they are the only fission yeast species. We do observe further cases of lineage-specific patterns in clades with distinct ecologies.

We found that nearly all examined polarization proteins were absent in the Microsporidia (Figure 2), including most of the conserved core. The only proteins that we observed are Iqg1 and Ubi4. Interestingly, we did not observe this pattern in the other basal phylum, the Cryptomycota. We believe that our observation is a true lineage-specific loss in the Microsporidia, as the majority of the polarization proteins (29 out of 34 proteins) are found in non-fungal eukaryotes, such as animals, amoeba and/or plants (see Supplementary File 5). The genomes of the parasitic Microsporidia are known to be highly condensed and lack other essential proteins, such as MAP kinases and proteins involved in stress response (Miranda-Saavedra et al. 2007; Peyretaillade et al. 2011; Vivarès et al. 2002). This strong reduction in the proteome is hypothesized as an adaptation to their parasitic life style. It is currently not understood which proteins play a role in polarized cell growth in this genus.

In contrast to the strong reduction in the Microsporidia, we observed lineage-specific gain of polarization proteins in the budding yeast species Saccharomycetaceae. Bem4, Gic1 and Gic2 are all restricted to this clade (Figure 2). Various causes can be involved. Genome-wide comparisons across the eukaryote tree have identified an increase in proteins domains in the lineage towards the Ascomycota (Zmasek & Godzik 2011). Furthermore, a whole genome duplication occurred in the *Saccharomyces* lineage after the divergence from the *Kluyveromyces* lineage, and has resulted in many duplicated genes (i.e., paralogs) and instances of accelerated evolution (Kellis et al. 2004; Wolfe & Shields 1997). Our results indicate that different processes have resulted in a myriad of lineage-specific patterns across the fungal tree.

### The conserved core of polarization; functional versus structural conservation

We observe a group of sixteen core proteins that are recurrently present in the vast majority of examined species (Figure 5). Interestingly, the 16 core proteins cover all functional groups from Cdc42 regulators and effectors to proteins involved in cytokinesis and exocytosis (Figure 1). Even though this group consists of the most prevalent polarization proteins, it does not represent the absolute minimal system needed for polarization. In fact, the majority of species does not have the full set of core proteins (i.e., the complete core is present in 66 out of the 200 strains/species), which can be seen as another indicator of high uniqueness of structural constitution of the polarization network across fungi. Different strains/species might achieve functional conservation of the core functions of the network by having different combinations of core proteins. In fact, we observed 12 or more core proteins (i.e., 75%) in 191 strains/species (i.e., 95,5%). These results suggest that functional conservation of the polarization network is high, but that structural conservation, in the sense of network composition, of the individual proteins varies across the fungal strains/species.

Various protein characteristics have been elucidated that are (in part) responsible for protein network conservation, such as position within the network, whether the proteins are essential and the number of interactions (Liu et al. 2015; Giaever et al. 2002). We observed high proportions of essential proteins (6 out of 7 essential proteins are core proteins) and short single domain proteins (5 out of 10) for the core proteins (Figure 2). Selection is thought to be strong on these classes of proteins, because of their crucial functions and long protein domains (Pál et al. 2006; Buljan & Bateman 2009). These functional characteristics are based on studies in *S. cerevisiae* and could be less relevant in other species. We did not find significant differences in the number of genetic and physical interactions between the conserved core proteins and the non-core proteins. Interestingly, Cdc42, Bem1, Cdc24 and Cla4 were among the conserved core and have the most interactions with the other proteins. These proteins also take central parts in the polarization network, as key regulator (Cdc42; Etienne-Manneville 2004; Park & Bi 2007; Johnson 1999), scaffolding for protein complex (Cdc24 and Bem1; Butty et al. 2002), and signal transducing (Cla4; Johnson 1999). At the same time, we do find a low number of physical and genetic interactions for the core proteins, Rdi1, Axl2, Rho3.Our results indicate that factors like if a protein is essential and the number of interactions, only partially explain the conservation of core proteins. We did find a significant effect of protein abundance on the conservation of core proteins. This observation supports the hypothesis that conserved proteins are generally more expressed in a cell, as discussed previously (Wall et al. 2005; Drummond 2005).

### Link to causal factors for variation in the polarization protein network

Here we aim to uncover potential causal factors influencing protein network evolution, in the sense of network size. Our MFA results show that the factors lifestyles, lineage and genetic distance co-vary with the size of the protein repertoire. These results indicate that the evolutionary background, adaptation to specific lifestyles (i.e., yeast-like, unicellular and filamentous) and evolutionary time, and thus an indirect measure of genetic drift, of a given species influence their polarization repertoire size. The examined factors do not explain all variation observed in the data, as 59% is undefined, indicative of missing causal factors. The discovery of, and interplay of, causal factors of adaptation and differentiation between species has gained much and long-term attention in the literature (Kimura 1967; Haldane 1927; Orr 2005; Masel 2011; Futuyma 2009). The long-lasting history of population genetics has shown that genetic variation, and thus sequence similarity and ultimately presence/absence of proteins, is caused by the interplay of mutation, natural selection, drift and gene flow, with descent and thus the heritable characteristics as the starting conditions. It is clear that not all these potential causal factors are incorporated in our study, mainly due to the scale of our study and the unavailability of the particular data for our focal species. Even though our analysis does not examine all factors that are likely to have played a role during the protein network evolution, we identified several factors that are, in part, responsible for the complex and highly dynamic polarization protein network evolution of fungi. Further expansion of experimental datasets and development of reliable large-scale comparative tools, should aid a better assessment of empirical data in light of the available theoretical models to study the full scope of real life protein network evolution.

Our study characterizes the fungal polarization protein network as highly dynamic across species, and we identify gene expression level, lineage, lifestyle and genetic drift, as factors correlating with the observed patterns of variation and adaptation. Our results provide further evidence that protein networks are often characterized by shared (ancient) conserved components as well as taxa-specific components that are variable between even closely related species. Our work sheds new light on the level and intensity of protein network evolution across broad and deep phylogenetic levels.

## Acknowledgements

We want to thank the members of the Laan Lab for providing critical feedback during earlier phases of this project. Thanks to W.KG. Daalman for advice on the statistical analyses. We would like to express our gratitude to the participants of the Gordon Research Conference on Cellular & Molecular Fungal Biology 2016 for constructive feedback on this project. Special thanks to Q. A. Justman, P. J. Boyton and E. M. Hyland for providing valuable comments on earlier drafts of this manuscript and to S.W.M. Pelders for assisting with writing python code. This work was supported by the Netherlands Organization for Scientific Research (NWO/OCW), as part of the Frontiers of Nanoscience program.

## Supplementary Files

Supplementary File 1. Proteins, Strains & Species information

Supplementary File 2. Protein Matrix Comparison 200 versus 36sp

Comparison of number of observed proteins between the full dataset of 200 strains/species and the 36 selected species with highest genome quality. Depicted are scatterplots for the following four lineages: Microsporidia (far left, squares), Saccharomycotina 1 representing the Saccharomycetaceae family (center left, circles), Saccharomycotina 2 representing e.g., the Debaryomycetaceae family (center right, triangles), Pezizomycotina (far right, diamonds). For each lineage a scatterplot of the full dataset (200 strains/species; right scatterplots) and the reduced dataset is depicted (36 species; left scatterplots). Lineages are color-coded as in Figure 4A. Medians (shown as grey lines) do not differ between the two datasets for all four lineages.

Supplementary File 3. Repeated Combinations Proteins

Supplementary File 4. MFA analysis

(A) Broken stick method. (B) Groups. (C) Correlation map of variables. (D) Partial axes.

Supplementary File 5. Eukaryote outgroups

